# Scaling Personalized TMS: A Scalp-Based MRI-guided Alternative to Neuronavigation

**DOI:** 10.1101/2025.09.05.673388

**Authors:** Farui Liu, Zong Zhang, Lijiang Wei, Yuanyuan Chen, Zeqing Zheng, Yilong Xu, Haosen Cai, Zheng Li, Yingying Tang, Jijun Wang, Chaozhe Zhu

**Author notes:** Address correspondence to: Chaozhe Zhu, Ph.D. State Key Lab of Cognitive Neuroscience and Learning & IDG/McGovern Institute for Brain Research, Beijing Normal University, Beijing 100875, China. These authors contributed equally to this work (Farui Liu, Zong Zhang).

## Abstract

**Background:** Personalized transcranial magnetic stimulation (TMS) has shown promise in treating psychiatric and neurological disorders, but its accessibility is limited by reliance on costly and complex neuronavigation systems, particularly in resource-constrained clinical settings.

**Objective:** To develop and validate a scalp-based alternative to neuronavigation for personalized TMS targeting.

**Methods:** We developed a Continuous Proportional Coordinate (CPC) system-based approach for personalized TMS targeting and validated its performance through three operators across ten participants, using individualized DLPFC-SGC targets as an example. The validation framework included targeting consistency assessment, reproducibility evaluation, and cross-cortical generalizability analysis.

**Results:** The CPC-based approach demonstrated comparable performance to neuronavigation, achieving 99.4% of its electric field strength (74.80 vs. 75.14 V/m; r = 0.962) and 93.9% of DLPFC-SGC functional connectivity (-0.216 vs. -0.230; r = 0.949), with high inter-operator reliability (ICC ≥ 0.903). Reproducibility analysis showed strong correlation between sessions (r = 0.948-0.957) with high reliability. Further simulations across 1,125 uniformly distributed cortical targets supported generalizability of this approach, with stable electric field preservation (99.07 ± 1.54%) across different brain regions.

**Conclusions:** The CPC-based approach offers a practical alternative to neuronavigation, maintaining targeting accuracy and providing a more accessible option for personalized TMS implementation.

## Introduction

Transcranial magnetic stimulation (TMS) is a non-invasive brain stimulation technique that generates electric fields in targeted cortical regions, modulating neural activity through both local and network effects [1,2]. While TMS has shown therapeutic potential in treating various psychiatric and neurological disorders, clinical outcomes remain variable [3]. Personalized TMS protocols, such as Stanford Accelerated Intelligent Neuromodulation Therapy (SAINT), have demonstrated significantly improved therapeutic efficacy, achieving response and remission rates of 80-90% in depression treatment, compared to conventional protocols which rarely exceed 50% [4]. Precise targeting of individualized cortical sites is a crucial determinant of therapeutic outcomes in TMS treatment. Currently, MRI-guided neuronavigation systems serve as the gold standard for implementing personalized TMS protocols [5]. These systems integrate patients’ physical space with their MRI data through anatomical landmark registration, enabling real-time tracking of coil position relative to the head [6]. Through visual feedback and error metrics, they facilitate precise stimulation of targeted sites, with their accuracy validated by both computational modeling and empirical studies [7,8].

Despite their precision, the widespread clinical adoption of neuronavigation systems remains limited by high costs and operational complexity [3, 9]. Recent surveys highlight this accessibility gap: only 7.4% of clinical TMS centers in Korea for stroke rehabilitation and 10% of Danish TMS facilities have neuronavigation capabilities [10,11]. This limited accessibility prevents personalized TMS protocols from reaching a broader patient population who could benefit from them, underscoring the urgent need for practical alternative targeting methods [3,12,13].

This study proposes and validates a Continuous Proportional Coordinate (CPC) system-based approach as a practical alternative to neuronavigation for personalized TMS targeting. The CPC system defines scalp positions using a pair of normalized coordinates (0-1), offering spatial mapping for MRI-derived cortical targeting with only a measuring tape needed for implementation (see Supplementary Fig. S1) [14–16]. We conducted comprehensive validation through three complementary components, using individualized DLPFC-SGC targets as an example. First, we compared the CPC-based approach with neuronavigation for targeting individualized DLPFC-SGC sites using ten 3D-printed head models derived from their MRI data. Three trained operators performed coil placements using both methods, and the induced electric field (EF) strengths and DLPFC-SGC functional connectivity (FC) were evaluated. Second, we assessed the reproducibility of the CPC-based method by conducting repeated coil placements across multiple sessions. Third, to demonstrate the method’s broad applicability across diverse brain regions, we performed computational simulations of coil placements and resulting electric fields at 1,125 cortical targets. The overall experimental design and CPC-based approach workflow are summarized in Figure 1.

**Figure 1.**
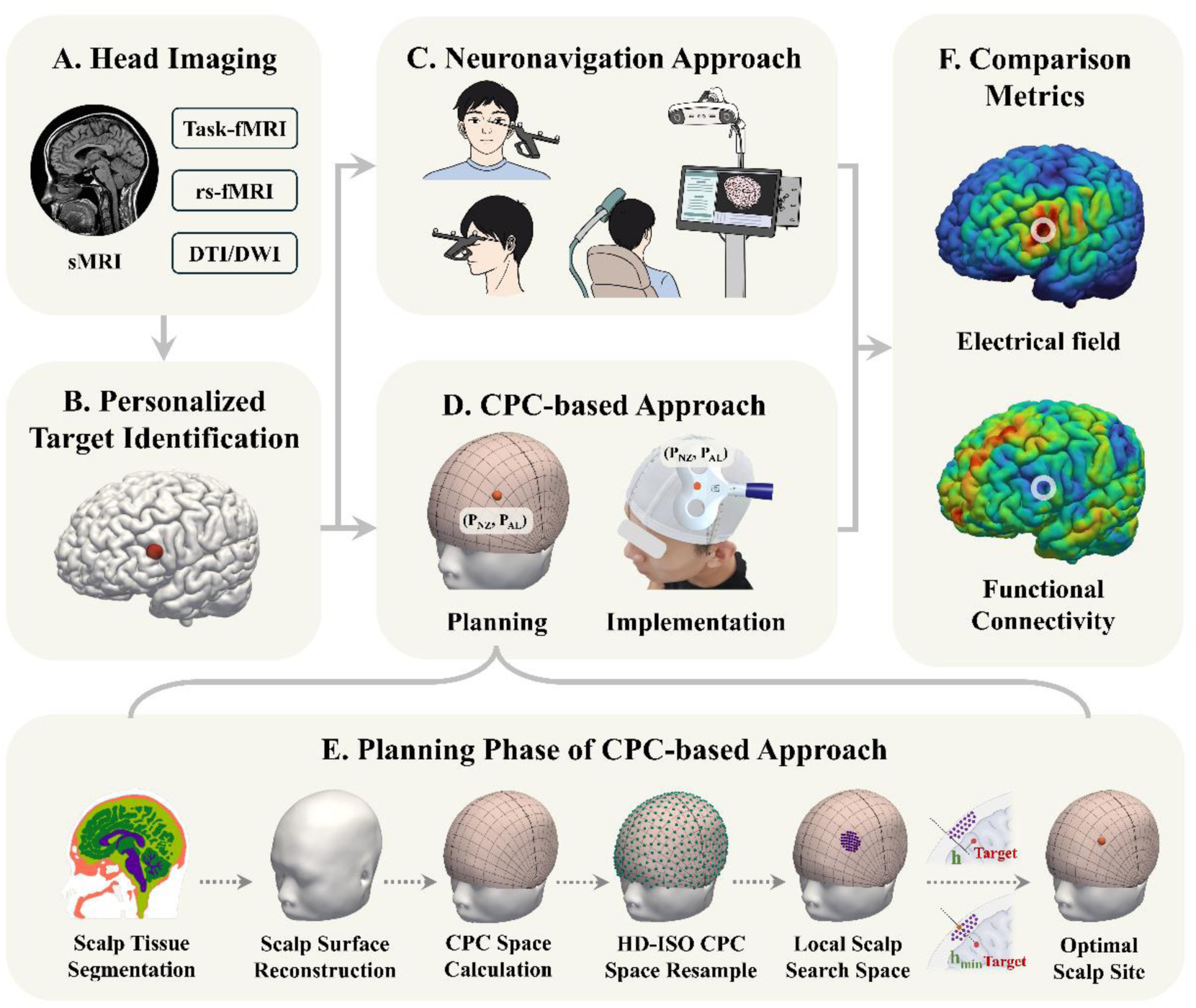
Overview of the CPC-based targeting approach and the consistency validation framework. (A-B) Target identification using multi-modal MRI data (structural MRI, task-based fMRI, resting-state fMRI, and DTI/DWI) to determine personalized targets. (C) Neuronavigation approach workflow including facial landmark registration, visual feedback, and coil placement. (D) The proposed CPC-based approach with planning and implementation phases. (E) Detailed workflow of the CPC-based planning phase, including: (1) scalp tissue segmentation and surface reconstruction, (2) CPC system calculation and HD-ISO resampling, (3) local scalp search space generation, and (4) optimal scalp site determination with CPC coordinates. (F) Validation framework comparing electric field strength and functional connectivity between neuronavigation and CPC-based approach.

## Method

### CPC-based MRI-guided Targeting Approach

The overall experimental design and CPC-based workflow are summarized in Figure 1. Figure 1A-B illustrates the process of individualized cortical target identification using multimodal MRI data (structural, task-based fMRI, resting-state fMRI, and DTI/DWI). Figure 1C depicts the conventional neuronavigation workflow involving facial landmark registration and real-time coil tracking. In contrast, the CPC-based approach (Fig. 1D) comprises two sequential phases: (1) a computational planning phase to determine optimal scalp coordinates for individual cortical targets, and (2) a practical implementation phase for manual coil placement. The detailed planning procedure is shown in Fig. 1E, including scalp surface reconstruction, CPC space generation, local search space definition, and optimization of the stimulation site. Here, we detail each phase of this approach.

### Planning Phase

The planning phase consists of sequential computational steps. First, scalp tissue is precisely segmented from individual structural MRI data and reconstructed into a 3D mesh surface using established computational tools (e.g., SimNIBS) [17]. Next, an individualized CPC scalp space is constructed based on the reconstructed surface, enabling standardized mapping of scalp locations. This CPC space is then resampled into high-density, isotropic (HD-ISO) scalp points with approximately 1 mm spacing using the Fibonacci series, ensuring precise spatial resolution for target mapping. For each specified cortical target, a local scalp search space is defined as a circular region with a 1 cm radius centered around its nearest scalp projection. Subsequently, an iterative optimization algorithm calculates perpendicular distances from the cortical target to the normal vectors of HD-ISO scalp points. This approach aligns with neuronavigation strategies by identifying the optimal stimulation site as the scalp point with minimal perpendicular distance to the target [16]. The final output comprises a pair of CPC coordinates corresponding to this optimal scalp position.

### Implementation Phase

Once the optimal scalp location is determined, coil placement involves a straightforward procedure requiring only a measuring tape. Operators first identify four standard anatomical landmarks (nasion, inion, and bilateral preauricular points) on the participant’s head. Using these landmarks as reference points, the target position is located by measuring proportional anterior-posterior and left-right distances based on the predetermined CPC coordinates. This systematic measuring approach enables precise marking of the stimulation site without specialized equipment. The TMS coil is then positioned with its center aligned to the marked location while maintaining tangential orientation to the scalp surface, following standardized protocols [15].

### Validation Experiment

#### Targeting Consistency Assessment

Ten participants (5 females and 5 males, mean age 19.9 ± 0.83 years) from the Southwest University Longitudinal Imaging Multimodal (SLIM) dataset were included in this study [18]. The dataset collection was approved by the Research Ethics Committee of the Brain Imaging Center of Southwest University on June 3, 2012, with the reference number spy-2012–012. The dataset provided high-resolution structural MRI (T1-weighted, 1mm isotropic) and resting-state functional MRI (8-minute duration) for each participant. T1-weighted anatomical images were acquired using a magnetization-prepared rapid gradient echo (MPRAGE) sequence. The parameters were as follows: repetition time (TR) = 1,900ms, echo time (TE) = 2.52ms, inversion time (TI) = 900ms, flip angle = 9°, resolution matrix = 256 × 256, number of slices = 176, slice thickness = 1.0 mm, and voxel size = 1 × 1 × 1 mm³. Resting-state functional magnetic resonance imaging (rs-fMRI) was conducted over an 8-minute period, during which participants were instructed to lie still with their eyes closed, remain awake, and refrain from focused thinking. A total of 242 contiguous whole-brain functional images were obtained using gradient-echo planar imaging (EPI) sequences. The imaging parameters were as follows: number of slices = 32, TR/TE = 2000/30 ms, flip angle = 90°, field of view (FOV) = 220 × 220 mm, slice thickness/gap = 3/1 mm, and voxel size = 3.4 × 3.4 × 3 mm³.

Individual DLPFC targets were identified using DLPFC-SGC FC, locating the left DLPFC region showing maximal anticorrelation with SGC based on resting-state fMRI analysis [19]. To facilitate experimental validation, we created 1:1 scale head models using participants’ reconstructed scalp surfaces, which were 3D-printed in epoxy resin with 50μm precision by a professional manufacturer. Detailed DLPFC target identification procedures and 3D printing specifications are provided in the supplementary materials.

To assesses targeting consistency, three trained operators conducted coil placement on each model using both the CPC-based method and neuronavigation (Brainsight, Rogue Research, Montreal, Canada). To prevent potential bias from prior knowledge, operators first performed CPC-based targeting before using neuronavigation. Coil position and orientation for both methods were recorded using the neuronavigation system for comparative analysis.

Targeting consistency was evaluated through EF simulations and FC analysis. EF simulations were conducted using SimNIBS 3.2.6 for each coil placement [17]. For the primary analysis, the region of interest (ROI) was defined as the intersection between a 5mm-radius sphere centered on each participant’s DLPFC-SGC target and their cortical mesh. Mean EF strength within the ROI was calculated for both methods. To examine the influence of ROI size on electric field calculations, additional analyses were performed using ROIs with different radii (3mm, 10mm, and 15mm). Using the SGC FC maps generated during target identification, DLPFC-SGC FC was computed at the actual stimulation sites for both methods, with stimulation sites determined using the 99.5% EF strength threshold.

The consistency between the methods was quantified using three complementary statistical approaches. First, Bland-Altman analysis assessed the agreement between methods by examining systematic differences and limits of agreement. Second, Pearson’s correlation coefficient evaluated the linear relationship between measurements from both methods. Finally, operator reliability was assessed using the intraclass correlation coefficient (ICC(2,1)), providing a measure of consistency across different operators.

#### Reproducibility Assessment

The reproducibility of the CPC-based method was assessed through a second coil placement session conducted four days after the initial measurements. The second session was performed by the three trained operators on the identical ten head models using the CPC-based method, following identical protocols from the consistency assessment. Coil positions and orientations were recorded using the neuronavigation system. EF simulations and FC calculations were performed primarily using the 5mm-radius ROI following procedures described in the consistency assessment. The influence of ROI size was also examined using additional radii (3mm, 10mm, and 15mm). Statistical analyses were performed using the metrics applied in the consistency assessment.

#### Cross-cortical Generalizability Assessment

To systematically evaluate the broad applicability of the CPC-based approach across diverse cortical regions, comprehensive computational simulations were conducted. As shown in Figure 3C, using a representative participant’s cortical mesh, a total of 1,125 targets were uniformly distributed across the cortical surface with approximately 8mm spacing. For each target, the region of interest (ROI) was defined as a 5mm-radius sphere centered on the target point. The optimal scalp position for each target was determined using the normal vector approach.

To simulate realistic targeting variations, we incorporated both positional and orientational deviations. For positional variations, we used the mean targeting error of 4.37mm observed in the consistency validation experiment, selecting six points at 60° intervals around each optimal position. For orientational variations, four coil orientations (0°, 45°, 90°, and 135°) were simulated at each deviated position. EF simulations were performed for all position-orientation combinations.

Targeting accuracy was quantified by calculating the ratio of mean EF strength within the ROI at deviated positions to that at the optimal position. For regional analysis, cortical regions were automatically segmented using volBrain system, and targeting accuracy was evaluated separately for frontal, temporal, parietal, and occipital regions [20].

## Results

### a) Targeting Consistency

Individual EF distributions for one operator are shown in Figure 2, covering all ten participants (S1-S10), with results from the other two operators provided in Fig. S2. The mean EF strength within the target ROI was 74.80 ± 11.50 V/m (mean ± SD) for the CPC-based method and 75.14 ± 10.13 V/m for neuronavigation, with the CPC-based approach reaching 99.36% of the neuronavigation values. Bland-Altman analysis revealed a mean difference of -0.34 V/m, with 95% limits of agreement ranging from - 6.75 to 6.07 V/m. The methods demonstrated strong correlation (r = 0.962, p < 0.001) and high operator reliability (CPC-based: ICC = 0.943, 95% CI: 0.85-0.98; Neuronavigation: ICC = 0.990, 95% CI: 0.97-1.00). These comparisons are illustrated in Figure 3A (left panel).

**Figure 2.**
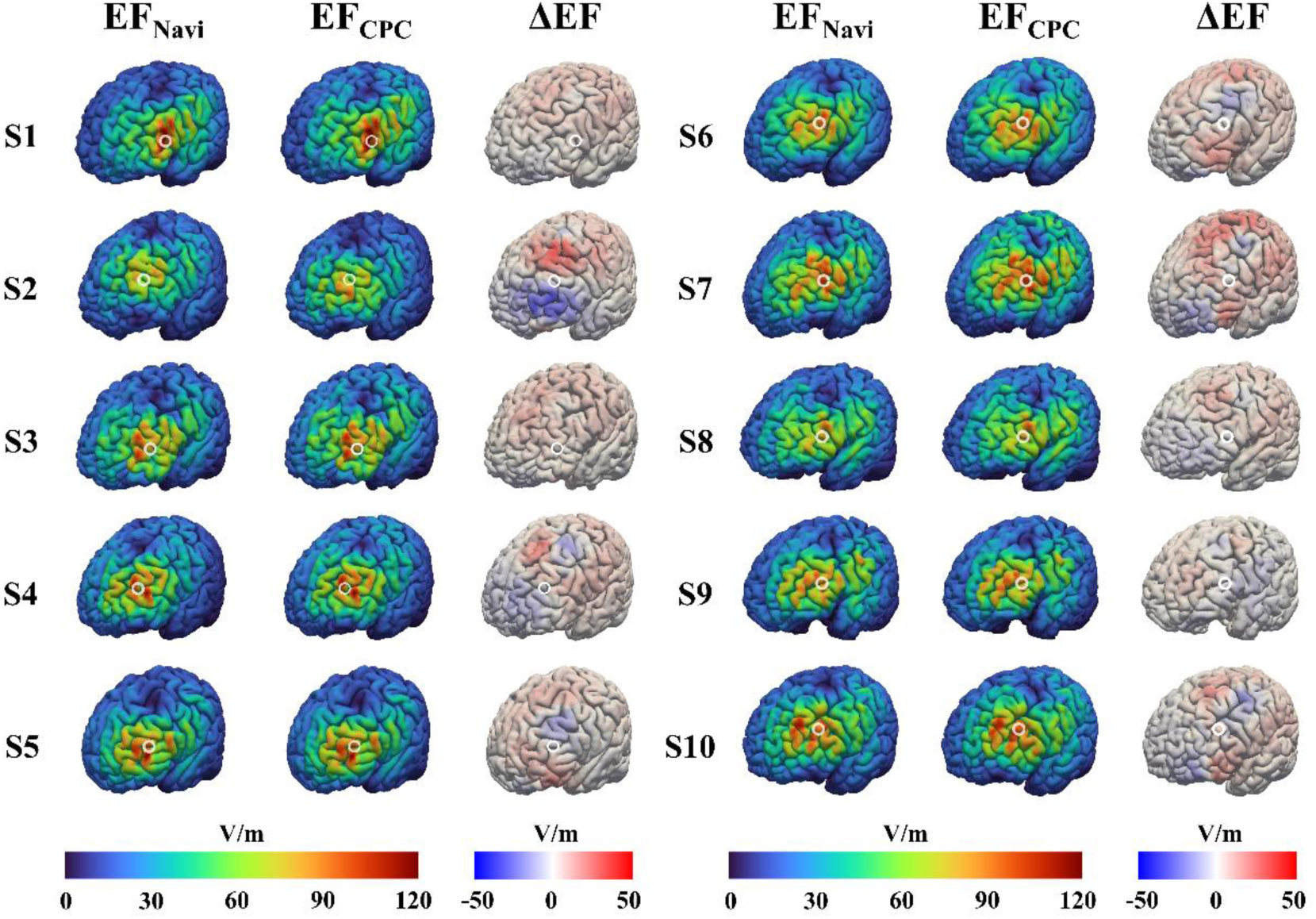
Comparison of electric field distributions between neuronavigation and CPC-based methods from operator 1. Electric field (EF) distributions induced by neuronavigation (EF_Navi) and CPC-based (EF_CPC) methods for ten participants (S1-S10), with their differences (ΔEF = EF_CPC - EF_Navi). The color bars indicate the electric field strength (V/m) for EF distributions (0-120 V/m) and differences (-50 to 50 V/m). Results from the other two operators are provided in Fig. S2.

For DLPFC-SGC FC analysis, the mean FC in the target ROI were -0.216 ± 0.080 (mean ± SD) for the CPC-based method and -0.230 ± 0.080 for neuronavigation, with the CPC-based approach achieving 93.90% of the neuronavigation values. Bland-Altman analysis showed a mean difference of 0.014, with 95% limits of agreement between -0.036 and 0.065. The methods demonstrated strong correlation (r = 0.949, p < 0.001) and high operator reliability (CPC-based: ICC = 0.903, 95% CI: 0.75-0.97; Neuronavigation: ICC = 0.993, 95% CI: 0.98-1.00). These results are shown in Figure 3A (right panel).

**Figure 3.**
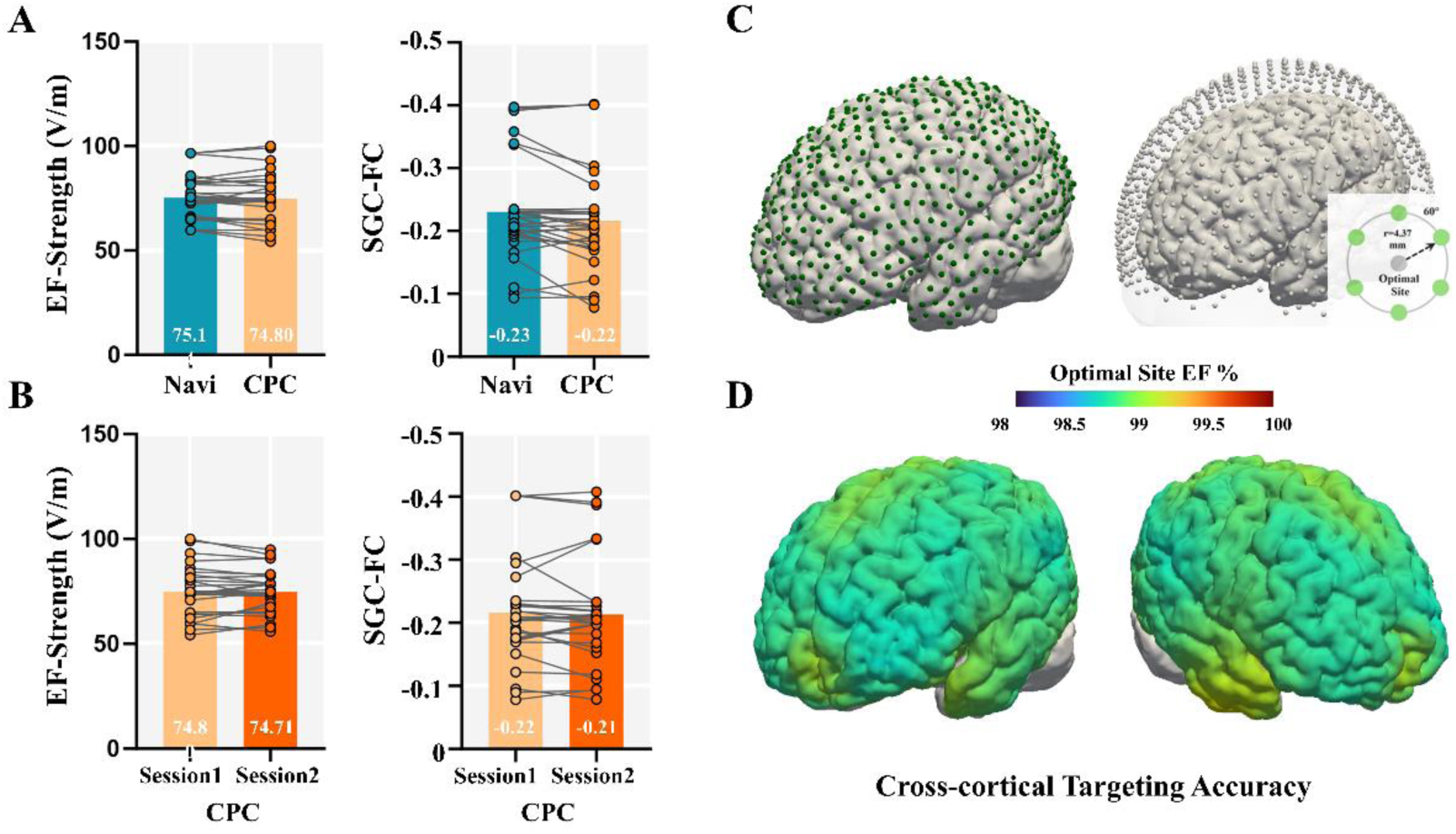
Validation of the CPC-based targeting approach. (A) Comparison between neuronavigation (Navi) and CPC-based methods for electric field (EF) strength and DLPFC-SGC functional connectivity (SGC-FC). (B) Test-retest reliability of the CPC-based method across two sessions for EF strength and SGC-FC measurements. Individual measurements are shown as connected dots and group means as bar plots. (C) Illustration of the cross-cortical generalizability simulation setup with uniformly distributed targets (8mm spacing, green dots) and deviated positions. (D) Regional analysis of EF preservation rates normalized to optimal targeting positions, with color scale indicating percentage of optimal EF strength (98-100%).

Additional analyses using different ROI sizes demonstrated consistent performance (Figure 4A). For the 3mm ROI, electric field strength values were CPC: 78.37 ± 13.92 V/m and Navigation: 78.84 ± 12.73 V/m, with strong correlation (r = 0.969, p < 0.001). Larger ROIs showed similar consistency, with 10mm ROI values of CPC: 65.63 ± 7.66 V/m and Navigation: 65.59 ± 6.15 V/m (r = 0.941, p < 0.001), and 15mm ROI values of CPC: 58.01 ± 6.09 V/m and Navigation: 57.71 ± 4.95 V/m (r = 0.929, p < 0.001). All ROI sizes maintained high operator reliability (ICC > 0.90) for both methods.

**Figure 4.**
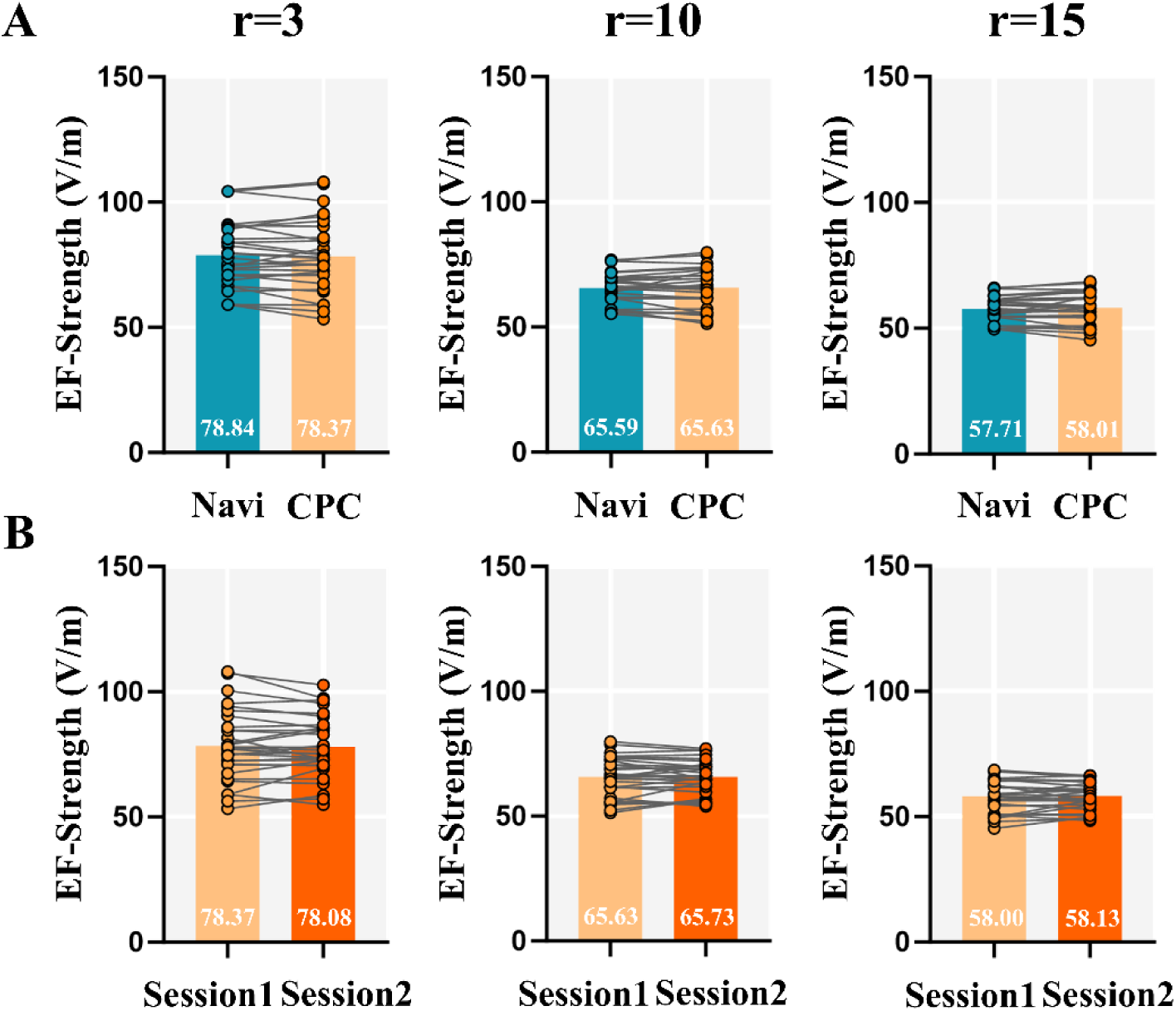
Analysis of ROI size effects on targeting performance. (A) Comparison between neuronavigation (Navi) and CPC-based methods using ROIs with radius of 3, 10, and 15 mm. (B) Test-retest reliability of the CPC-based method across two sessions for different ROI sizes. Individual measurements are shown as connected dots and group means as bar plots (in V/m).

### b) Reproducibility

EF strength across sessions demonstrated high consistency (Session 1: 74.80 ± 11.50 V/m, mean ± SD; Session 2: 74.71 ± 9.85 V/m). Bland-Altman analysis revealed a mean difference of 0.09 V/m with 95% limits of agreement between -7.35 and 7.54 V/m. The sessions showed strong correlation (r = 0.948, p < 0.001) and high operator reliability (Session 1: ICC = 0.943, 95% CI: 0.85-0.98; Session 2: ICC = 0.939, 95% CI: 0.84-0.98). These comparisons are illustrated in Figure 3B (left panel).

DLPFC-SGC FC measurements also showed consistent values (Session 1: -0.216± 0.080, mean ± SD; Session 2: -0.213 ± 0.083), with Session 2 measurements reaching 98.82% of Session 1 values. Bland-Altman analysis revealed a mean difference of - 0.003, with 95% limits of agreement between -0.050 and 0.044. The sessions demonstrated strong correlation (r = 0.957, p < 0.001) and high operator reliability (Session 1: ICC = 0.903, 95% CI: 0.75-0.97; Session 2: ICC = 0.869, 95% CI: 0.68-0.96). These results are shown in Figure 3B (right panel).

Test-retest reliability remained robust across different ROI sizes (Figure 4B). For the 3mm ROI, between-session measurements were Session 1: 78.37 ± 13.92 V/m and Session 2: 78.08 ± 11.99 V/m (r = 0.956, p < 0.001). Similar reliability patterns were observed for 10mm ROI (Session 1: 65.63 ± 7.66 V/m, Session 2: 65.73 ± 6.64 V/m; r = 0.913, p < 0.001) and 15mm ROI (Session 1: 58.01 ± 6.09 V/m, Session 2: 58.13 ± 5.19 V/m; r = 0.894, p < 0.001). All ROI sizes maintained high operator reliability (ICC > 0.90) for both sessions.

### c) Cross-cortical Generalizability

As illustrated in Figure 3D, simulation results across 1,125 cortical targets showed stable EF preservation relative to optimal positions, with a mean preservation rate of 99.07 ± 1.54% (mean ± SD). Analysis across different coil orientations revealed consistent preservation (0°: 99.05 ± 1.17%; 45°: 99.11 ± 2.86%; 90°: 99.04 ± 0.96%; 135°: 99.11 ± 2.86%). Regional analysis demonstrated comparable preservation rates for frontal (99.02 ± 0.35%, range: 97.82-100.88%), temporal (99.46 ± 3.14%, range: 91.01-146.67%), parietal (98.90 ± 0.15%, range: 98.59-99.58%), and occipital regions (98.89 ± 0.16%, range: 98.61-99.47%).

## Discussion

This study proposes a CPC-based alternative for implementing personalized TMS targeting when neuronavigation is not accessible. Our validation experiments demonstrated that the CPC-based method offers a practical and cost-effective solution for clinical settings. The results show strong consistency between the CPC-based and neuronavigation approach, as evidenced by high ICC for both induced EF strength and functional connectivity at individualized SGC-DLPFC targets. Furthermore, test-retest assessment across different sessions confirmed the method’s robust reproducibility. Computational simulations across multiple cortical targets further demonstrated the method’s performance remained consistent across different brain regions, with electric field characteristics preserved despite typical positioning variations.

The CPC-based approach also offers practical advantages for clinical implementation. By requiring only basic measuring tools and straightforward procedures, it enhances operational efficiency for both operators and patients. While conventional neuronavigation procedures typically require 20-30 minutes for setup and registration [21], our validation experiment demonstrated that the CPC-based approach can be completed within 4.4 minutes while maintaining comparable targeting accuracy [15]. These findings suggest that the CPC-based approach could facilitate broader clinical adoption of personalized TMS protocols, particularly in resource-limited settings where neuronavigation systems are not be readily available, by balancing targeting accuracy with practical efficiency.

Several previous studies have explored alternative approaches to neuronavigation for personalized TMS targeting. For example, triangulation-based methods have been developed to locate specific targets such as the motor cortex and Broca’s area using individual structural MRI [12,22]. However, these methods were typically designed for specific cortical targets, making their anatomical reference points target-dependent and potentially arbitrary, without establishing a generalized framework applicable to arbitrary cortical targets. Some approaches even required additional markers during MRI acquisition, introducing procedural complexity [23]. Notably, most of these methods lack systematic evaluation of their targeting performance across diverse cortical regions.

Unlike neuronavigation systems, which provide real-time tracking and feedback for position adjustment and head movement compensation, our method cannot monitor coil position and orientation during stimulation [24]. Potential technical solutions, such as implementing a positioning block with corresponding slots on the coil for enhanced stability and orientation control, could address this limitation, though such modifications would require systematic validation in future studies.

Furthermore, this study utilized high-resolution structural MRI data from a single center. The impact of MRI quality parameters (including resolution, signal-to-noise ratio, and acquisition time) on targeting accuracy requires systematic investigation, particularly regarding the reliability of scalp surface reconstruction across different imaging protocols and scanner types [25].

In recent years, cerebellar TMS has gained increasing attention in both clinical treatment and cognitive research [26]. However, the CPC system has inherent geometric limitations in covering certain anatomical regions, particularly over the cerebellum, which may affect its applicability for cerebellar stimulation protocols.

The current study employs a simplified geometric model for TMS targeting, similar to conventional neuronavigation systems. However, to better account for the complexities of TMS-induced electric fields, future developments could incorporate more sophisticated modeling approaches, potentially integrating machine learning techniques for real-time electric field estimation [27,28]. This aligns with our previously proposed Scalp Geometry-based Parameter (SGP) system, where the combination of optimal scalp positioning and coil angle optimization could enable more precise targeting protocols [15]. While some studies have explored 3D-printed positioning solutions, these approaches face challenges including material safety concerns, coil support design complexity, and patient tolerability issues [29,30].

Most importantly, large-scale clinical trials are needed to validate the therapeutic effectiveness of this approach and establish its clinical utility in real-world settings. These future developments could help bridge the gap between advanced personalized TMS protocols and their widespread clinical implementation, potentially improving treatment accessibility and outcomes for a broader patient population.

## Acknowledgements

This work was granted support from the National Natural Science Foundation of China (Grant No. 82071999 and No. 61431002) and the Lingang Laboratory (Grant No. LG-TKN-202205-01).

## Ethics Statement

The Southwest University Longitudinal Imaging Multimodal (SLIM) dataset used in this study was approved by the Research Ethics Committee of the Brain Imaging Center of Southwest University on June 3, 2012, with the reference number spy-2012–012. The privacy rights of human subjects have been observed, and informed consent was obtained from all participants.

## Data and Code Availability Statement

The datasets and analysis code supporting the findings of this study are available from the corresponding author upon reasonable request, subject to the following conditions: (1) execution of an appropriate data sharing agreement to ensure compliance with privacy regulations and institutional policies; (2) approval from the requesting researcher’s institutional review board or ethics committee, where applicable; (3) submission of a detailed research proposal outlining the intended use of the data and methodology; and (4) potential requirement for collaborative arrangement or co-authorship depending on the extent of data utilization and contribution to derivative works. These requirements are implemented to ensure responsible data sharing while protecting participant privacy and maintaining research integrity.

