## Supplementary for "Scaling Personalized TMS: A Scalp-Based MRI-guided Alternative to Neuronavigation"

### CPC system

The previously proposed Continuous Proportional Coordinate (CPC) system is used to represent individual scalp locations in this study [1,2]. The CPC system, defined based on five cranial landmarks (the nasion (NZ), the inion (IZ), the left and right preauricular points (AL/AR), and the vertex (CZ)), is a continuous coordinate system that enables a one-to-one mapping of each point on the scalp to pNZ and pAL coordinates (Fig. S1). This is critical for the precise mapping of personalized TMS cortical targets onto the scalp. Leveraging these properties, the recently proposed Scalp Geometry-based Parameter (SGP) space - a derivation of the CPC system for the purpose of describing TMS coil positions - has demonstrated good performance in manual localization of coils spatially and temporally [3].

For those interested in implementing this technique, a comprehensive tutorial demonstrating the complete process of manual coil placement is available at https://www.youtube.com/watch?v=8nMs_IaWwUg. This visual guide provides step-by-step instructions, offering practical insights into the application of the CPC and SGP systems in TMS targeting.

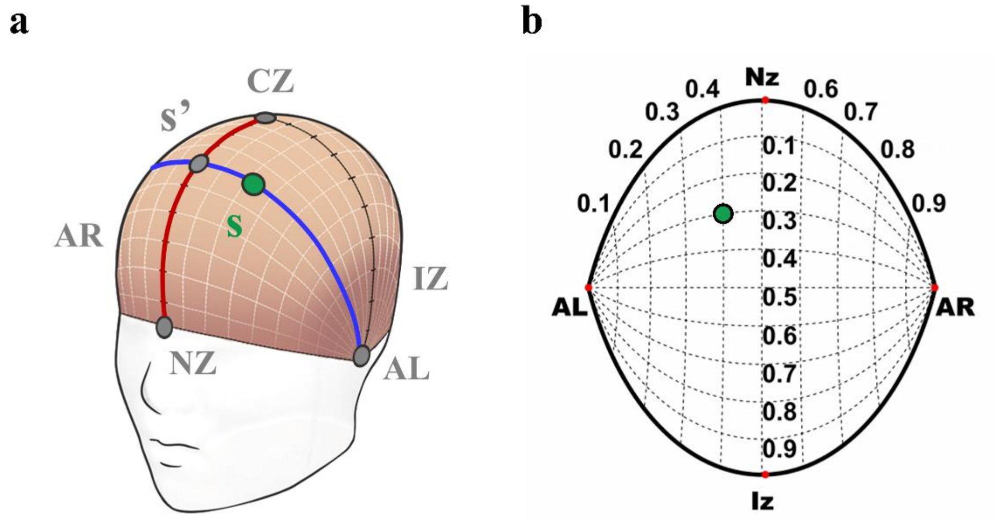


**Figure S1. CPC system definition**. (a) Any location on the scalp can be represented using a pair of proportional coordinates ($p_{NZ}$, $p_{AL}$) within the range of 0 to 1, expressing the location’s anterior-posterior and left-right proportions, respectively. Specifically, a reference curve (in red) is defined as the intersection of the scalp surface with the mid-sagittal plane through NZ, CZ, and IZ. For a specific scalp location $\boldsymbol{s}$ (green point), an active curve (in blue) is delineated, traversing points AL, AR, and $\boldsymbol{s}$, and intersecting the reference curve at point $\boldsymbol{s}'$ (gray point). The CPC coordinates of $\boldsymbol{s}$ are calculated as follows: $p_{NZ} = L_{NZ-s'}/L_r$ and $p_{AL} = L_{AL-s}/L_a$. Here, $L_{NZ-s'}$ is the length from NZ to s’ along the reference curve, and Lr is the full length of the reference curve. $L_{AL-s}$ is the length from AL to $\boldsymbol{s}$ along the active curve, and La is the full length of the active curve. (b) The 2D Hammer-Aitoff projection of the CPC system.

#### Image preprocessing

Functional MRI data preprocessing was performed using SPM12 software, following the protocol described by Fox et al. (2012) [5]. The initial four volumes were discarded to allow for magnetic field stabilization. Subsequent volumes underwent slice timing correction and realignment. Functional images were co-registered with individual T1 images and normalized to the Montreal Neurological Institute (MNI) space using a unified segmentation algorithm. The images were then resampled to 2-mm isotropic voxels and smoothed using a 6-mm full-width at half-maximum (FWHM) Gaussian kernel. Additional preprocessing steps, including low-pass filtering ($f < 0.08$ Hz) and removal of nuisance covariates (head motion parameters, white matter signal, cerebrospinal fluid signal, and global signal), were conducted using the DPABI toolbox (Data Processing & Analysis for Brain Imaging).

#### Individual SGC-DLPFC Target Identification

For each participant, functional connectivity (FC) between the DLPFC target region and subgenual cingulate cortex (SGC) was calculated. The DLPFC was delineated using 20-mm radius spheres centered on three specific locations in the left hemisphere: Brodmann area 9 [MNI coordinates: −36, 39, 43], Brodmann area 46 [MNI coordinates: −44, 40, 29], and a TMS site located 5 cm from the prefrontal cortex [MNI coordinates: −41, 16, 54] [6,7]. The SGC was defined as a 10-mm radius sphere centered at MNI coordinates [6, 16, -10], encompassing only gray matter. This location represents the average position where reductions in SGC activity have been observed across various effective antidepressant treatments [5].

The DLPFC time series were obtained by averaging the time series of all gray matter voxels within the DLPFC ROI. Due to the relatively low signal-to-noise ratio in the SGC, its time series was computed using an SGC-based seed map [5,6]. Specifically, the SGC-based seed map represented the connectivity of the SGC ROI to all gray matter voxels in the whole brain, averaged across a group of 114 healthy individuals [1,2]. This map was used as a weighting factor to compute the weighted mean of the time series of all gray matter voxels in each participant (excluding those within the DLPFC and SGC ROIs). The resulting weighted mean time series was employed as the SGC time series.

Pearson correlation coefficients were calculated between the time series of the DLPFC and SGC for each participant, and subsequently transformed into z-scores using Fisher's z transformation. The target was selected as the center of a sphere of 6 mm radius exhibiting the strongest anti-correlation with the SGC [6]. Each sphere was weighted by its proximity to the cortical surface to mimic the linear decay of the TMS field.

#### Individual Optimal Scalp Site Determination

The T1-weighted images of the ten participants were processed using SimNIBS 3.2.6 for tissue segmentation and individual scalp surface reconstruction. To enhance the quality of the reconstructed scalp surfaces, a 10-step Laplacian smoothing algorithm was applied [8]. Subsequently, the CPC system was constructed based on these smoothed scalp surfaces and further refined through resampling into high-density, 1mm isotropic scalp points (approximately 72,000 points) using the Fibonacci series. To efficiently locate the optimal stimulation site, a local search space was defined, comprising the 700 scalp points nearest to each individual's SGC-DLPFC target. This number of points corresponds to an approximate circular area with a radius of 1 cm on the scalp surface, providing a focused yet sufficiently broad search region. Within this confined area, an iterative optimization process utilizing a normal vector model was employed to determine the optimal scalp location and its corresponding CPC coordinates for each participant's individual SGC-DLPFC target [2].

#### Scalp Surface Reconstruction and 3D-Print

For each individual, head tissue segmentation and initial surface reconstruction were performed using the *headreco* command in SimNIBS 3.2.6. To enhance the quality of the reconstructed scalp surfaces, a 10-step Laplacian smoothing algorithm was subsequently applied using MeshLab. The surface quality of the digital models was significantly improved by this smoothing process. After optimization, the refined models were sent to a specialized 3D printing company. Physical models were produced by the company using epoxy resin, with a precision of 50μm and at a 1:1 scale. Highly accurate physical representations of the subjects' head structures were created through this approach, which combined computational modeling techniques with advanced 3D printing.

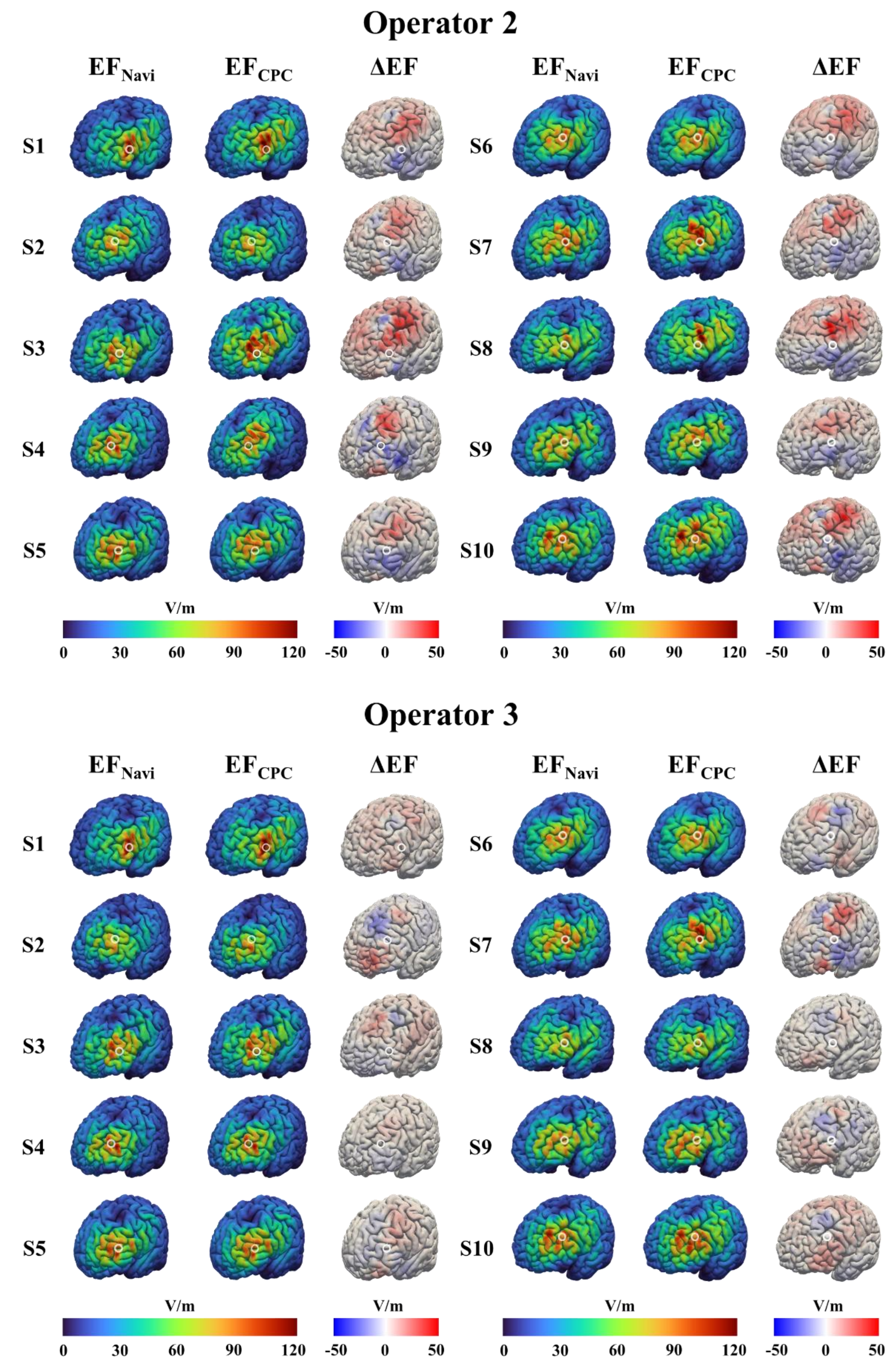


**Figure S2. Validation results of the CPC-based targeting approach from two additional operators.** Electric field (EF) distributions induced by neuronavigation ($EF_{Navi}$) and CPC-based ($EF_{CPC}$) targeting methods for all ten participants (S1-S10), along with their differences (ΔEF).

### Supplementary Analysis of Individual Operator Performance

For electric field strength analysis, individual operator results demonstrated consistent performance. With the CPC method, mean values were 74.80 ± 12.3 V/m, 73.70 ± 10.8 V/m, and 75.09 ± 11.9 V/m for Operators 1, 2, and 3, respectively. Corresponding neuronavigation values were 75.14 ± 11.2 V/m, 75.36 ± 10.5 V/m, and 74.92 ± 10.1 V/m. Similarly, functional connectivity measures showed consistency across operators. CPC-based measurements yielded -0.226 ± 0.074, -0.200 ± 0.092, and -0.222 ± 0.078 for Operators 1, 2, and 3, respectively. Neuronavigation measurements were -0.231 ± 0.082, -0.229 ± 0.085, and -0.231 ± 0.075. Individual operator data are visualized in Figure S3.

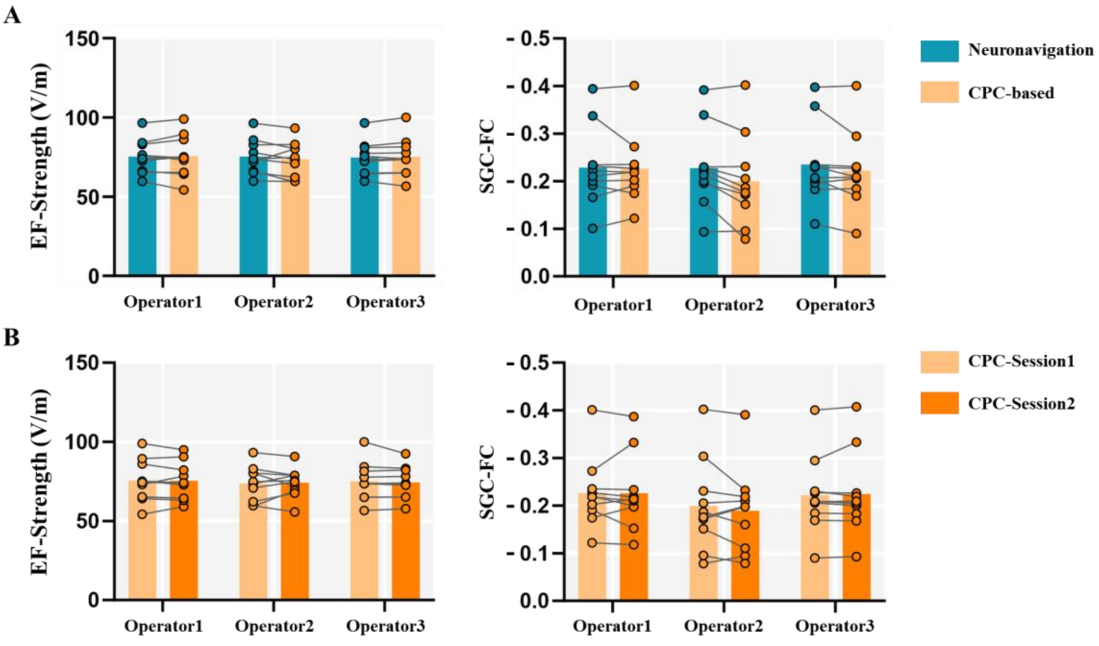


**Figure S3. Individual operator performance in coil targeting consistency and t** **reliability.** Left panels show electric field strength, right panels show DLPFC-SGC f connectivity. A) Comparison between navigation and CPC method for each operator retest comparison of CPC method across sessions for each operator.
